# Orthoformimycin inhibits translation elongation by displacing the A-site tRNA and preventing peptide bond formation

**DOI:** 10.1101/2021.11.29.470217

**Authors:** Andreas Schedlbauer, Tatsuya Kaminishi, Attilio Fabbretti, Pohl Milón, Xu Han, Borja Ochoa-Lizarralde, Retina Çapuni, Claudio O. Gualerzi, Sean R. Connell, Paola Fucini

**Affiliations:** Center for Cooperative Research in Biosciences (CIC bioGUNE), Basque Research and Technology Alliance (BRTA), Bizkaia Technology Park, Building 801A, 48160 Derio, Spain; Graduate School of Medicine, Osaka University, 2-2 Yamadaoka, Suita, Osaka, 565-0871 Japan; Laboratory of Genetics, Department of Biosciences and Veterinary Medicine, University of Camerino, 62032 Camerino, Italy; Centre for Research and Innovation, Faculty of Health Sciences, Universidad Peruana de Ciencias Aplicadas (UPC), Lima, Peru; IKERBASQUE, Basque Foundation for Science, 48011 Bilbao, Spain; Structural Biology Unit, BioCruces Bizkaia Health Research Institute, Plaza Cruces s/n, 48903, Barakaldo, Spain; Research Centre for Experimental Marine Biology and Biotechnology (PiE-UPV/EHU), University of the Basque Country (UPV/EHU), Areatza z/g, 48620 Plentzia-Bizkaia, Basque Country, Spain

## Abstract

The ribosome is a major target for antibiotics owing to its essential cellular role in protein synthesis. Structural analysis of ribosome-antibiotic complexes provides insight into the molecular basis for their inhibitory action and highlights possible avenues to improve their potential or overcome existing resistance mechanisms. Here we use X-ray crystallography and pre-steady state kinetics to detail the inhibitory mechanism of the antimicrobial on the large ribosomal subunit.

## INTRODUCTION

The World Health Organisation (WHO) cautions that antimicrobial resistance (AMR) is leading to bacterial infections that are more difficult to treat while the EU estimates that AMR costs €1.5 billion per year in healthcare costs and productivity losses (*1*, *2*). While there is no singular solution to this growing health issue it is evident that new antimicrobial agents are needed. Natural product isolation has been historically one way to discover antimicrobials but one could say these don’t represent novel antibiotics as they are well-known to nature and thus AMR mechanisms (*3*). That said these natural scaffolds are important as they inherently possess chemical properties suitable for antibiotics and thus serve as starting point or inspiration for the development of antibiotics (*3*, *4*). Key to exploiting these natural products is characterizing them from a structure/function perspective. Orthoformimycin (ofm) is one such natural product from *Streptomyces* that inhibits protein synthesis. It was recently isolated in a high throughput screen of 6700 microbial fermentation extracts (*5*). Initial characterization indicated ofm is a potential antimicrobial with modest but measurable activity *in vivo*, being most effective against *Moraxella catarrhalis* (MIC = 32 μg mL^-1^) (*5*). Importantly, *in vitro* protein synthesis assays revealed ofm was active against bacterial systems but not yeast-based translation systems indicating bacterial specificity. Functionally ofm was shown to inhibit protein synthesis suggesting it targets the ribosome while *in vitro* assays suggested it interacts with the large subunit, inhibiting peptide bond formation and EF-G associated activities (*5*).

Structurally ofm is composed of three moieties: a [4-(dimethyl-amino)-phenyl]-2-methylprop-2-enoic acid (DMPE; cinnamic-like moiety), a aminocyclitol, and L-erythronic acid (*5*) (**Figure 1A**). A DMPE-like moiety is also found in three other potent inhibitors of protein synthesis, puromycin (pmn), hygromycin A (hgr), and A201A, while a similar cyclitol part is also present in hgr (*6*, *7*). These three antibiotics all bind within the A site (*6*, *7*) of the 50S ribosomal subunit, in a cleft that is solely comprised of 23S rRNA and includes residues from the peptidyl transferase center (PTC). The PTC is highly conserved and is the active centre on the 50S subunit that binds the aminoacyl (A)- and peptidyl (P)-tRNAs to catalyse peptide bond formation between their aminoacyl substrates. When pmn, hgr and A201A bind within the A-site cleft they sterically overlap with the 3’ terminus of the A-tRNA (*6*, *7*). The DPME-like moiety of pmn harbours an amino group at the C2 position equivalent to the aminoacyl-tRNA allowing it to accept the amino acid or nascent chain attached to the P-tRNA and thus pmn can participate in the peptide bond formation reaction. In contrast hgr and A201A lack an acceptor group in their chemical scaffolds because the 2-amino seen in pmn is substituted with a methyl group and accordingly hgr and A201A are unable to serve as active substrates for the PTC. Thus, while pmn, a tyrosyl-tRNA-mimic, inhibits translation by being irreversibly incorporated into the C-terminus of the nascent polypeptide chain attached to the P-tRNA and promoting its premature release (*8*), hgr and A201A use steric hindrance to interfere with the binding of ligands to the A-site (*6*, *7*).

**Figure 1:**
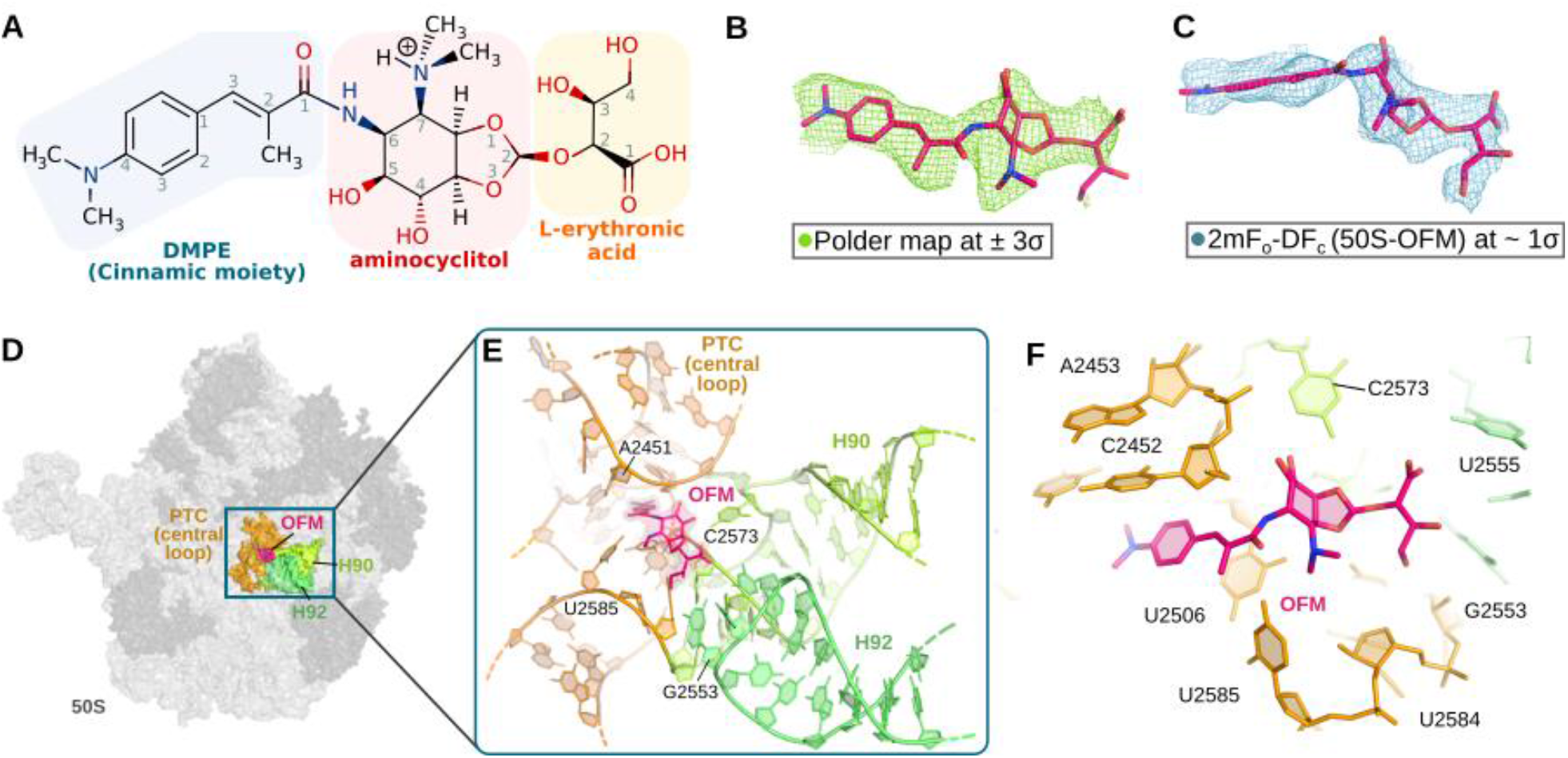
Chemical structure, electron density maps and ofm binding pocket on the 50S ribosomal subunit. (**A**) Chemical structure of ofm. (**B**) Unbiased Polder map and (**C**) *2Fo-Fc* electron density maps of ofm in complex with the 50S ribosomal subunit (green and blue mesh, respectively). Additional views and the conventional mFo-DFc initial unbiased difference map are shown in **Supplemental Figure 1A-D**, while the complete *2Fo-Fc* electron density maps for the ligand binding pocket are shown **in Supplemental Figure 1E-F.** (**D**) The overview of the drug binding pocket on the 50S subunit with the refined model, at a scale highlighting the rRNA helices (**E**) and the rRNA residues (**F**) forming the ofm binding pocket. The geometric centres of panels E and F are rotated approximately 90° with respect to each other. Note while U2506 is modelled with a dual conformation only the higher occupancy conformation is shown in the figures for clarity but both conformations are compatible with ligand binding. Throughout the figures and text *E. coli* numbering is used instead of *D. radiodurans* for clarity.

Following previous reports (*5*) that ofm binds the large ribosomal subunit like the structurally similar pmn, hgr and A201A, we used *in vitro* functional assays and X-ray crystallography to localize and describe the ribosomal binding site of ofm. A comparison with chemically similar antimicrobials reveals a range of interactions that contribute to the binding mode of A-site targeting antibiotics.

## RESULTS

### Overall Structure and binding pocket interactions

To determine the ofm binding site, vacant 50S subunit crystals from *Deinococcus radiodurans* were soaked with 20 μM ofm and X-ray crystallography was used to solve the structure of the complex refined at 3.0 Å. The initial unbiased difference Fourier map (mFo-DFc) determined using phases derived from the deposited *apo 50S* subunit (PDBID 2ZJR (*9*)) showed only weak fragmented density attributed to the drug in the A site of the PTC. However, extensive improvement of the search model for the 50S subunit, substantially improved the quality of the initial unbiased map allowing a confident placement of the drug (**Figure 1B**) within the ribosomal A site. Further refinement of the 50S-ofm model led to a 3.0 Å resolution structure with the map for ofm displaying clear quasi planarity of the DMPE moiety with the conjugated peptide bond-like part (**Table 1; Figure 1C**).

**Table 1.** Summary of Crystallographic Data collection and Refinement statistics.

| Data collection | 50S-OFM | 50S-apo |
| --- | --- | --- |
| Space group | I222 | I222 |
| Cell dimensions (a,b,c) [Å] | 170.460, 409.370,<br>697.800 | 169.90, 410.76,<br>696.12 |
| Resolution range | 59.45 -2.80 (2.95-2.80) | 59.3–2.90 (3.06–2.90) |
| Resolution where $I/\sigma I=2$ [Å] | 3.0 | 3.2 |
| # of observed reflection | 14234761(1445878) | 11 664 835 (1 754 909) |
| # of unique reflections | 591820 (85103) | 531 662 (78 466) |
| Multiplicity | 24.1 (24.6) | 21.9 (22.3) |
| Completeness [%] | 99.9 (99.8) | 99.8 (99.6) |
| $R_{\text{merge}}$ [%] | 25.2 (744.9) | 24.1 (786.2) |
| Mean $I/\sigma(I)$ | 9.88 (0.89) | 10.38 (0.71) |
| CC(1/2) [%] | 99.9 (49.5) | 99.9 (47.8) |
| Wilson B-factor [Å<sup>2</sup>] | 81.9 | 78.94 |
| **Refinement statistics** |  |  |
| Rwork/Rfree [%] | 24.5/27.6 | 24.6/28.1 |
| **Model composition** |  |  |
| Chains | 31 | 31 |
| Non-hydrogen atoms | 88350 | 88322 |
| Protein residues | 3255 | 3255 |
| Nucleotides | 2929 | 2929 |
| Ligands | 1 | 0 |
| Zn | 1 | 1 |
| Mg | 252 | 302 |
| **RMSD deviations from ideal values** |  |  |
| Bond length (Å) | 0.017 | 0.020 |
| Bond angles (°) | 2.708 | 2.979 |
| **Ramachandran plot (%)** |  |  |
| Favored | 95.16 | 95.28 |
| Allowed | 4.81 | 4.72 |
| Outliers | 0.03 | 0.00 |
| **Other structural quality metrics** |  |  |
| MolProbity score | 2.13 | 2.26 |
| Clash score | 13.2 | 19.5 |
| Rotamer outliers (%) | 1.64 | 1.53 |
| Cβ outliers (%) | 1.16 | 1.23 |
| Non-trans peptide planes (%) | 0 | 0 |
| Cis proline | 0 | 0 |
| Twisted proline | 0 | 0 |

Values in parentheses refer to the highest resolution bin. The data was obtained from a single crystal.

The ofm binding pocket is formed by 23S rRNA residues of the highly conserved peptidyl transferase central loop and helices 90 (H90) and 92 (H92; A-loop; **Figure 1D-F**) which is consistent with chemical probing experiments **(Supplemental Figure 2)**. In the back of the pocket (near the ribosomal tunnel), the planar DMPE moiety is the most buried part of ofm as it inserts into a crevice normally occupied by the aminoacyl moiety of the A-tRNA and is sandwiched between C2452 and U2506 of the 23S rRNA (**Figure 2A**). In this position the 4-(dimethyl-amino)-phenyl is involved in a parallel shifted aromatic π stack with the base of C2452, whereas the alkene group of the drug stabilizes a trans U2506-U2585 non-Watson-Crick base pair interaction by an alkene-arene π stacking interaction with the base of residue U2506 (**Figure 2A,B**). In fact, when the structures of the apo 50S subunit and the 50S-ofm complex are compared, it is evident that ofm also induces conformational changes in the PTC including displacement of a ribosome bound magnesium and a slight displacement in U2506 and U2585 (**Figure 2C**) allowing a non-Watson-Crick base pair to form (**Figure 2B,C and Supplemental Figure 3**). This U2506-U2585 interaction is also observed in the presence of A-site ligand hgr (*6*, *7*) but the rearrangement is distinct from that induced by the proper binding of an A-site substrate (*10*) indicating that ofm, although binding the A-site, does not activate it like a canonical substrate.

**Figure 2.**
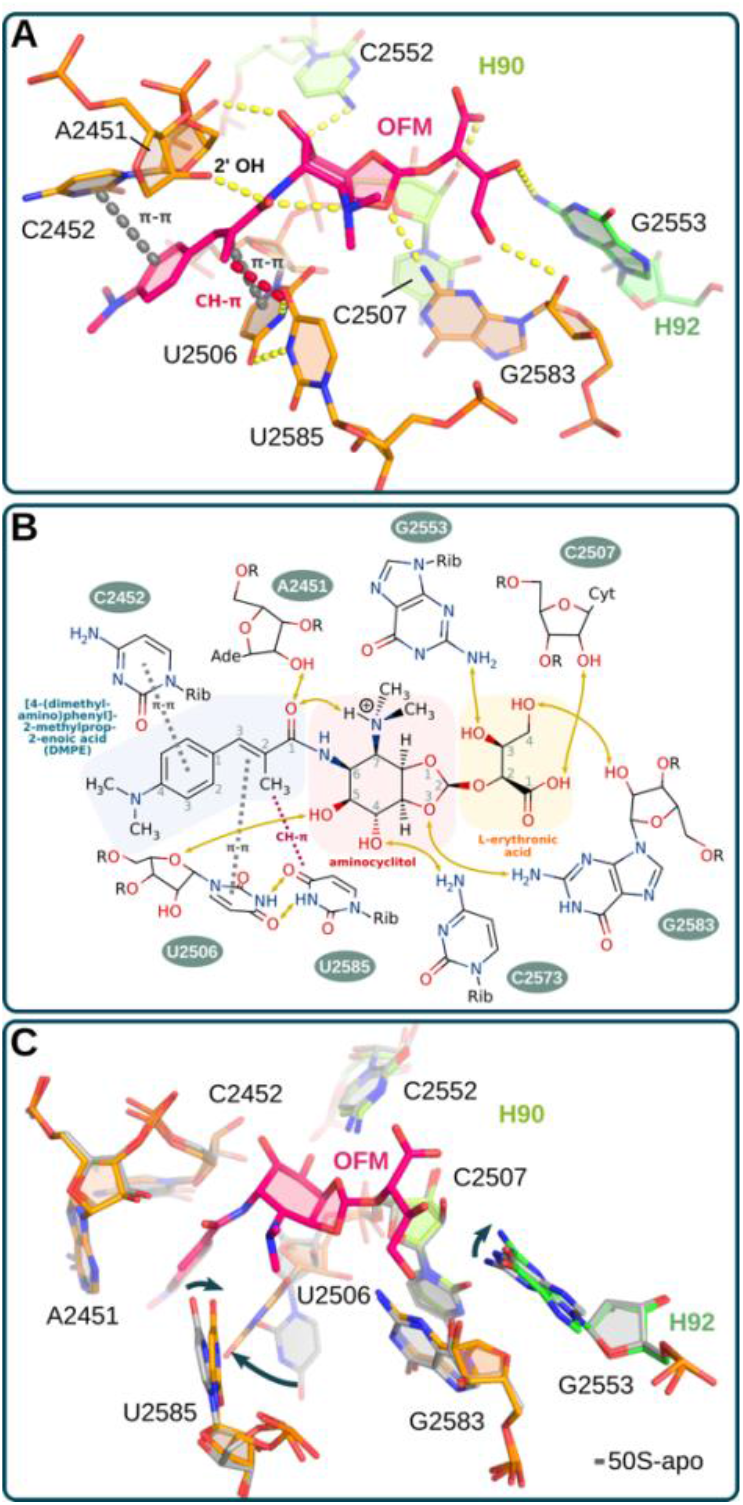
Interactions and conformational changes within the ofm binding site. Putative interactions based on geometric criteria between ofm and nucleotides of the 23S rRNA forming the binding pocket residues are shown on (**A**) the 3D model and (**B**) schematically in a 2D representation. Aromatic ring and alkene-arene π stacking interactions are colored grey, CH-π in red, and hydrogen bonds are drawn in yellow. (**C**) Binding of ofm to the A-site side of the PTC induces conformational changes in 23S rRNA residues (U2506, G2553 and U2585) involved in A-site tRNA binding.

The central aminocyclitol moiety is more exposed with one face open to the intersubunit space and is bound in the crevice normally occupied by the backbone/ribose of A76 in the accommodated A-tRNA. Here the extra-cyclic hydroxyl groups at positions C4 and C5 are within hydrogen bonding distance of C2573 and U2506 in the PTC, respectively, while O3 could hydrogen bond with G2583 (**Figure 2A,B**). The C7 dimethyl amine is chemically distinct from the hydroxyl group normally found in a similar position of the other A-site binding drugs (A201A, pmn, and hgr). This group introduces a positive charge and a hydrophobic surface near U2585 and A2602 and can form an intermolecular polar hydrogen bond with carbonyl oxygen O1 of DMPE moiety to stabilize the conformation of the ligand (**Figure 2A,B**). Finally, the erythronic moiety, which extends ofm towards the A-loop, can hydrogen bond with G2553 (**Figure 2A,B**) resulting in its displacement relative to that seen in the 50S-apo structure (**Figure 2C**). Normally C75 of the accommodated A-tRNA base pairs with G2553 (*11*, *12*) and thus, by filling this space the erythronic moiety of ofm can block this interaction with the tRNA. Overall, the ofm structure indicates that, like the chemically similar A-site targeting antibiotics, ofm functions by binding within the A-site cleft in a position normally occupied by the 3’ CCA-end of the A-tRNA.

#### Orthoformimycin prevents correct positioning of the aminoacyl-tRNA

As ofm binds within the cleft used by the 3’ CCA-end of the A-tRNA during protein synthesis we predict ofm would interfere with peptide bond formation. Accordingly, we assayed the ability of the A-site-bound aminoacyl-tRNA to act as an acceptor substrate in peptide bond formation in the presence and absence of ofm. As seen in **Figure 3A** and **B (red line)** both the kinetics and steady state levels of transpeptidation are strongly inhibited by increasing concentrations of ofm. For example, an IC50 of just 3±0.2 μM in the assay measuring fMet-Phe-tRNA formation (**Figure 3B**; red line) and a IC50 of 1.6±0.1 μM in the *in vitro* translation system that assays all reactions of the elongation phase in aggregate (**Figure 3B**; black line). These steady state values are higher than those published for hgr (*7*) while the kinetic experiments (**Figure 3A)** indicate ofm is a very effective inhibitor of peptide bond formation.

**Figure 3.**
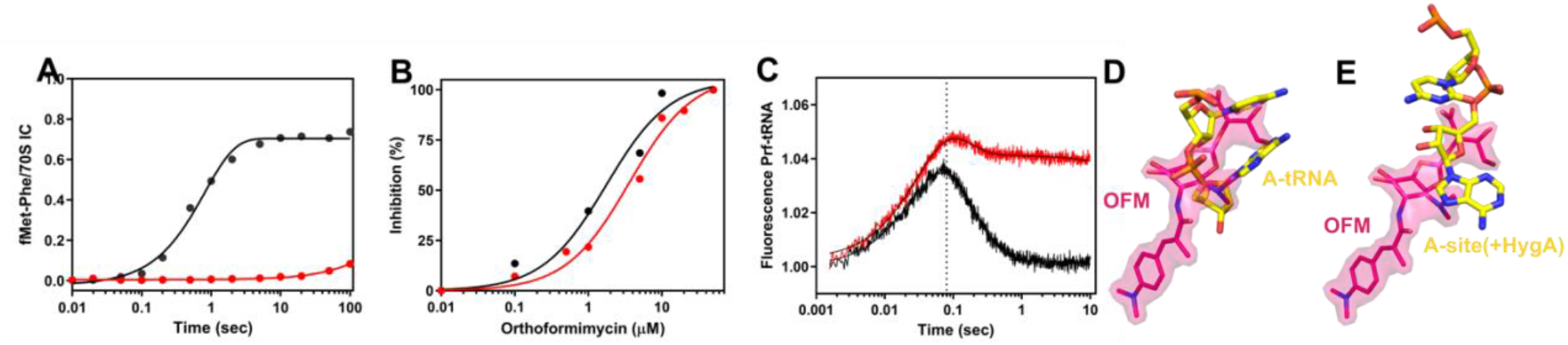
Effect of orthoformimycin on A-site binding of aminoacyl-tRNA and on peptide bond formation. (**A**) Effect of the orthoformimycin on the kinetics of transpeptidation as measured by quenched flow. (**B**) orthoformimycin concentration dependence of the levels of mRNA translation (black) and transpeptidation (red) inhibition. (**C**) Kinetics of EF-Tu-dependent A-site binding of proflavine-labelled Phe-tRNA^Phe^ to mRNA-programmed 70S ribosomes carrying P-site bound fMet-tRNA^fMet^ (70S *IC*) in the presence (red tracing) and absence (black tracing) of 100 μM orthoformimycin. The binding kinetics was followed in a fluorescence stopped-flow apparatus as described in Materials and Methods. (D-E) The steric clash between ofm and a fully accommodated A-tRNA and a partially accommodated A-tRNA (stalled by hgr) is illustrated by aligning the ofm-50S structure with PDBs (fully accommodated A-tRNA:7K00; partially accommodated A-tRNA:5DOY).

The ability of the A-site substrate to participate in transpeptidation is dependent on the A-tRNA being correctly accommodated onto the A-site, a multi-step process that occurs first on the 30S (decoding site) and then on the 50S (PTC) subunit. Ofm does not appear to effect tRNA binding and decoding on the 30S subunit, both in the presence or in the absence of EF-Tu (fluorescence increase in **Figure 3C**)(*5*), a finding in-line with the fact that ofm binds to the 50S and not to the 30S ribosomal subunit. Consistently, kinetic analysis of EF-Tu-dependent binding of Phe-tRNA to the 70S ribosome using fluorescence stopped-flow showed an impairment only in the final adjustment of the tRNA, which is attributed to GTP hydrolysis, EF-Tu:GDP dissociation, and/or Phe-tRNA accommodation (fluorescence decrease in **Figure 3C**). This finding indicates that an ofm may alter the placement of the acceptor end of the A-tRNA in the peptidyl transferase center of the 50S. As described above, ofm sterically clashes with the acceptor end (C75/A76) of the fully accommodated A-tRNA (**Figure 3D**). This overlap with the 3’end of the A-tRNA is evident when superimposing the 50S-ofm model with a structure showing an accommodated A-tRNA bound to the 50S subunit, which shows that the DMPE moiety overlaps the aminoacyl moiety of the A-tRNA, the aminocyclitol and L-erythronic groups clash with the accommodated A-tRNA near A76 and C75, respectively (**Figure 3D**). Wilson and colleagues (*6*) have also determined a structure of a partially accommodated A-tRNA that is blocked from entering the A-site cleft by the chemically similar antibiotic hgr. A structural alignment with this accommodating A-tRNA stalled by hgr shows that the L- erythronic group, which is not present in hgr, still clashes with this intermediate A-tRNA (**Figure 3E**), suggesting that ofm could block the A-tRNA earlier in the accommodation process than hgr. This could explain why ofm is an effective inhibitor of transpeptidation in kinetic experiments. A puzzling effect of orthoformimycin is the occurrence of EF-G-dependent aminoacyl-tRNA translocation apparently uncoupled from mRNA movement (*5*). The incorrect position acquired by A-site aminoacyl-tRNA in the presence of the antibiotic is probably at the root of this effect. Despite the failure of accepting an amino acid from P-site-bound tRNA, incoming EF-G recognizes the complex, hydrolyzes GTP and induces the movement of the A-site aminoacyl-tRNA. However, the movement of this tRNA, non-correctly adjusted in the decoding site of the 30S subunit fails to transmit the movement to the mRNA that remains immobile. The slower dissociation of the Pi from EF-G (*5*) is the likely consequence of this pseudo-translocation.

### OFM as a potent structural template for antibiotic-antibiotic conjugates

The binding site of ofm is shared by the A-tRNA and other chemically similar A-site targeting antibiotics (i.e., pmn, hgr and A201A) although each has their own mode of interaction/action owing to their individual chemical structure **(Figure 4)**. Common to all these ligands is an aromatic amino acid-like moiety that binds within the cleft formed by C2452 and U2506 of the 23S rRNA. However, the A-site targeting antibiotics are structurally more diverse when compared in the regions that extend into the ribosomal tunnel or into the A-site cleft (**Figure 4**). For example, ofm harbours an additional subunit, the L-erythronic moiety (yellow), that extends the scaffold towards and interacts with the A-loop (**Figure 4**). This subunit potentially makes additional hydrogen bonds with the 23S rRNA pocket that are not possible with either hgr or A201A (**Figure 2B**) and more strongly clashes with the A-tRNA (**Figure 3D,E**). In contrast, in the ribosomal tunnel, ofm has the shortest extension, while hgr and A201A have 1 or 2 sugar subunits appended at the C4 position, respectively (**Figure 4**). In the case of hgr, the furanose moiety is critical for antimicrobial activity (*13*) and in the case of both hgr and A201A these sugars interact (hydrogen bond) with the ribosome possibly contributing to their affinity (*6*). Importantly, the sugars also extend into the binding sites for the macrolide antibiotics and thus hgr and A201A bind competitively with the macrolides to the ribosome (*6*, *14*). Specifically, A201A competes with both the 14-membered macrolide erythromycin and 16-membered macrolides like tylosin, while hgr competes with the latter only (*6*, *14*). As seen in **Figure 5A-B,** the shorter dimethyl amine extension of ofm does not reach into the macrolide binding site and thus opens the possibility that using ofm or designing compounds with ofm-like extensions could be used effectively in combination with the macrolide class of antibiotics. Additionally, in **Figure 5A-B,** it can be seen that the macrolide antibiotics tylosin and erythromycin are oriented relative to ofm for the design of novel antibiotic-antibiotic conjugates. The sugar subunit of hgr is also assisted with resistance mechanisms. For example, the self-resistance mechanism, Hyg21 (*15*), employed by the hgr producing *S. hygroscopicus*, involves phosphorylating the 2’’-OH of the furanose and therefore lack of this sugar subunit would prevent Hyg21 from conferring cross resistance to the chemically similar ofm. Polikanov *et al*. (*6*) have also shown that the Cfr resistance mechanism, which methylates A2503 to confer resistance to several peptidyl transferase inhibitors (chloramphenicol, lincomycin and linezolid), is also active against hgr as the furanose subunit would clash with a m^8^A2503 modification. Modelling this methylation in the ofm-50S structure (**Figure 5C)** suggest that the shorter dimethylamine modification in ofm would not clash with the m^8^A2503 modification and thus, we predict ofm would not be sensitive to the Cfr resistance mechanism. Together we hypothesize that the ofm dimethylamine modification could represent a minimal C4 extension that preserves antimicrobial activity, but importantly could bypass existing resistance mechanisms active against the chemically similar hgr.

**Figure 4.**
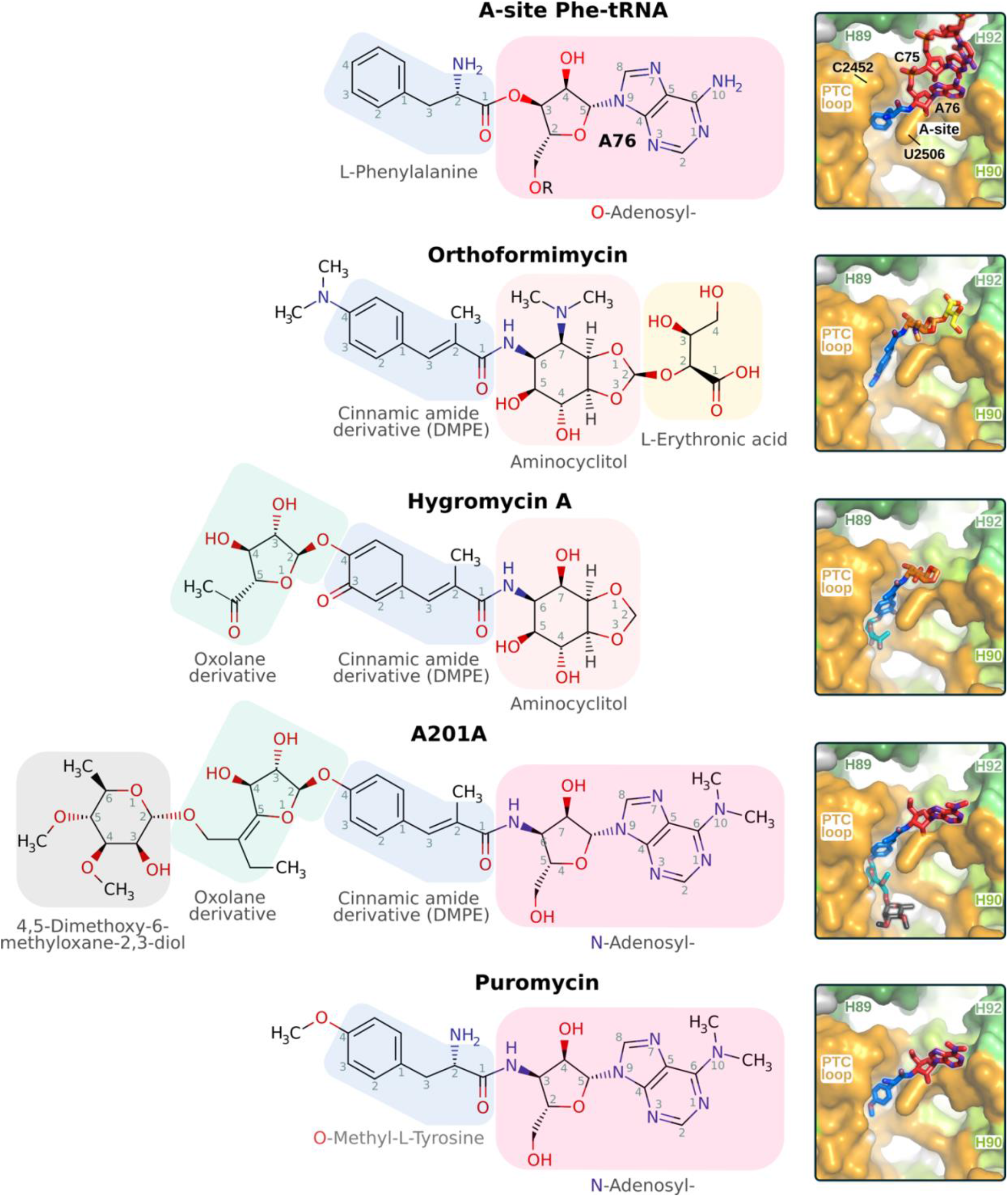
Similarities of ribosomal A-site binders based on their chemical scaffold composition. (Left) Comparison of the chemical scaffold of ligands specifically interfering with the full accommodation of the A site tRNA at the 50S ribosomal subunit in comparison with the structure of an A-site Phe-tRNA (bottom). The applied color code reflects the similarity of the chemical building units. (Right) Structural accommodation of the A-site tRNA with C75 and A76 labeled (top) and of the ligands at their ribosome binding pockets.

**Figure 5.**
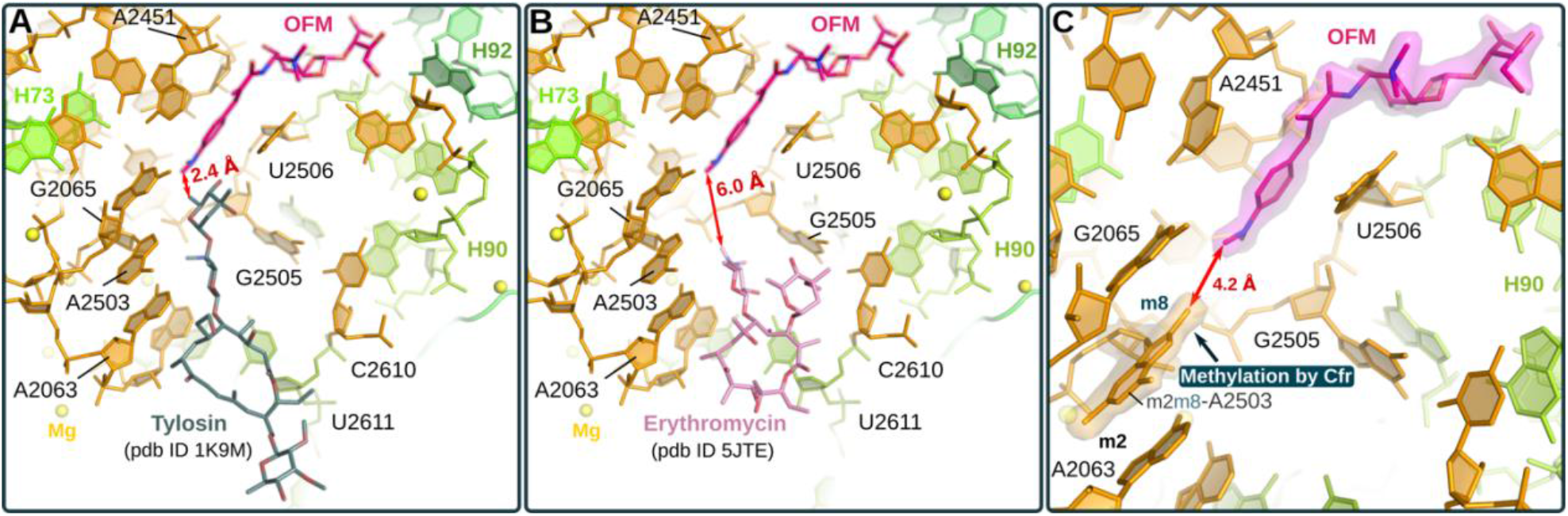
OFM as a potent structural template for the creation of fusion antibiotics and its binding conformation in the OFM/dr50S complex suggests a possible avenue overcome of the Cfr resistance mechanism effective against other antibiotics targeting the ribosomal A-site. (**A**,**B**) The binding modes of the macrolide antibiotics tylosin and erythromycin in their ribosomal 50S complexes (aligned on the OFM/dr50S complex based on residues of the 23S rRNA located within 30Å of the OFM binding site) indicate their proper orientation relative to OFM for the design of novel antibiotic-antibiotic conjugates (AACs). (**C**) Based on distance analysis OFM binding appears to be not impaired by the presence of the modification introduced by Cfr, a radical SAM methylase that methylates the base of residue A2503 at the C8 position (next to the methylation at C2 present in wild type ribosome of *Escherichia coli*). This modification has been identified to be sufficient for causing resistance to phenicols, lincosamides, oxazolidinones, pleuromutilins, streptogramin A, A201A, and hygromycin A.

## DISCUSSION

The structure presented here indicates that ofm binds within the A-site, sterically clashing with the fully accommodated A-tRNA. Accordingly, we posit that this steric clash will lead to it functioning like hgr and A201A (*6*) and cause the accommodating A-tRNA to oscillate between intermediate states (i.e., between the A/T-like conformation and partially accommodated state) thus stalling translation. Finally, our structural analysis of ofm expands our understanding of the interaction mode used by A-site targeting antibiotics and highlights several avenues that could be used to modify the common pmn, hgr, ofm, and A201A scaffolds to develop more effective anti-microbials that bypass existing resistance mechanisms.

## ACKNOWLEDGMENTS

This work could not have been performed without the expert assistance of the staff at the X06SA beam line (Swiss Light Source [SLS], Villigen). This work was supported by grants from Bizkaia:Talent and the European Union’s Seventh Framework Program (Marie Curie Actions to S.C., A.S., T.K.), the Marie Curie Actions Career Integration Grant (PCIG14-GA-2013-632072 to P.F.), the Ministerio de Economía Y Competitividad Grant (MINECO; CTQ201782222-R to P.F. and S.R.C.), the *Fondo Nacional de Desarrollo Científico, Tecnológico y de Innovación Tecnológica* (154-2017-FONDECYT to P.M.), and FIRB Futuro in Ricerca from the Ministero Italiano dell’Istruzione, dell’Università e della Ricerca (RBFR130VS5 001) to A.F. We thank MINECO for the Severo Ochoa Excellence Accreditation (SEV-2016-0644).

## EXPERIMENTAL PROCEDURES

### X-ray crystallography

Crystals of the 50S ribosomal subunit from *Deinococcus radiodurans* were prepared by the vapor diffusion method in sitting drops at 19°C as described previously (*16*, *17*) and soaked overnight in a stabilizing solution containing 20 μM orthoformimycin (ofm) prior to flash-freezing in liquid nitrogen. X-ray diffraction data were recorded at the beamline X06SA of the Swiss Light Source (Villigen, Switzerland) equipped with a PILATUS 6M pixel detector, and processed to 2.8 Å using the XDS (*18*) and CCP4 (*19*) software packages. As a search model for molecular replacement a previously published apo structure of the *D. radiodurans* 50S subunit (PDB accession code: 2ZJR) was used and refined against the newly collected 50S-OFM data in Phenix (*20*) followed by iterative rounds of manual model building in COOT (*21*) and ISOLDE (*22*). Prior to refinement about 5% of the experimental observations were removed from the reflection data for the free R factor calculation. The binding location and orientation of the drug as well as structural alteration of the 23S rRNA close to the binding site were unambiguously determined from unbiased initial mF_obs_–DF_model_ difference and Polder omit maps contoured at +/-3σ (**Supplemental Figure 1**). The 4-(dimethyl-amino)-phenyl moiety being structurally in conjugation with the alkene of 2-methylprop-2-enoic acid part (see **Figure 2**) were restrained during refinement to be in-plane to each other (omitting the two methyl groups attached to the amine due to partial sp^3^ character of the amine nitrogen). Although the molecular structure of the ligand indicates a preference for a further continuous planarity encompassing the peptide bond like moiety to the aminocyclitol part its carbonyl and amide atoms (of the peptide bond) weren’t restrained to maintain strict planarity taking the empirical analyses of atomic-resolution protein structures (*23*) into account. No torsion restraints were applied along the aminocyclitol ring dihedrals enabling optimization into any chair like or twist boat conformation. Starting coordinates for the drug were created using Openbabel 6.0 (http://openbabel.org/wiki/)](*24*) and restraint information about its geometry were derived using eLBOW (*25*). The structural models were visualized by PyMOL (The PyMOL Molecular Graphics System, Version 2.5.0, Schrödinger, LLC). Numbering from *Escherichia coli* was used for residues of the rRNA throughout this manuscript.

### Biochemical assays

*In vitro* mRNA translation (driven by 027IF2Cp(A) mRNA) and dipeptide (fMet-Phe) formation were carried out as described (*26*, *27*). *In situ* probing of the 23S rRNA by hydroxyl radical cleavage was carried out as described (*28*). EF-Tu-dependent A-site binding of proflavine-labeled Phe-tRNA to MFmRNA-programmed 70S ribosomes carrying P-site bound fMet-tRNA (70S *IC*), was carried out in the absence or presence of 100 μM ofm as described (*29*).

