## Supplemental Figures for "Orthoformimycin inhibits translation elongation by displacing the A-site tRNA and preventing peptide bond formation"

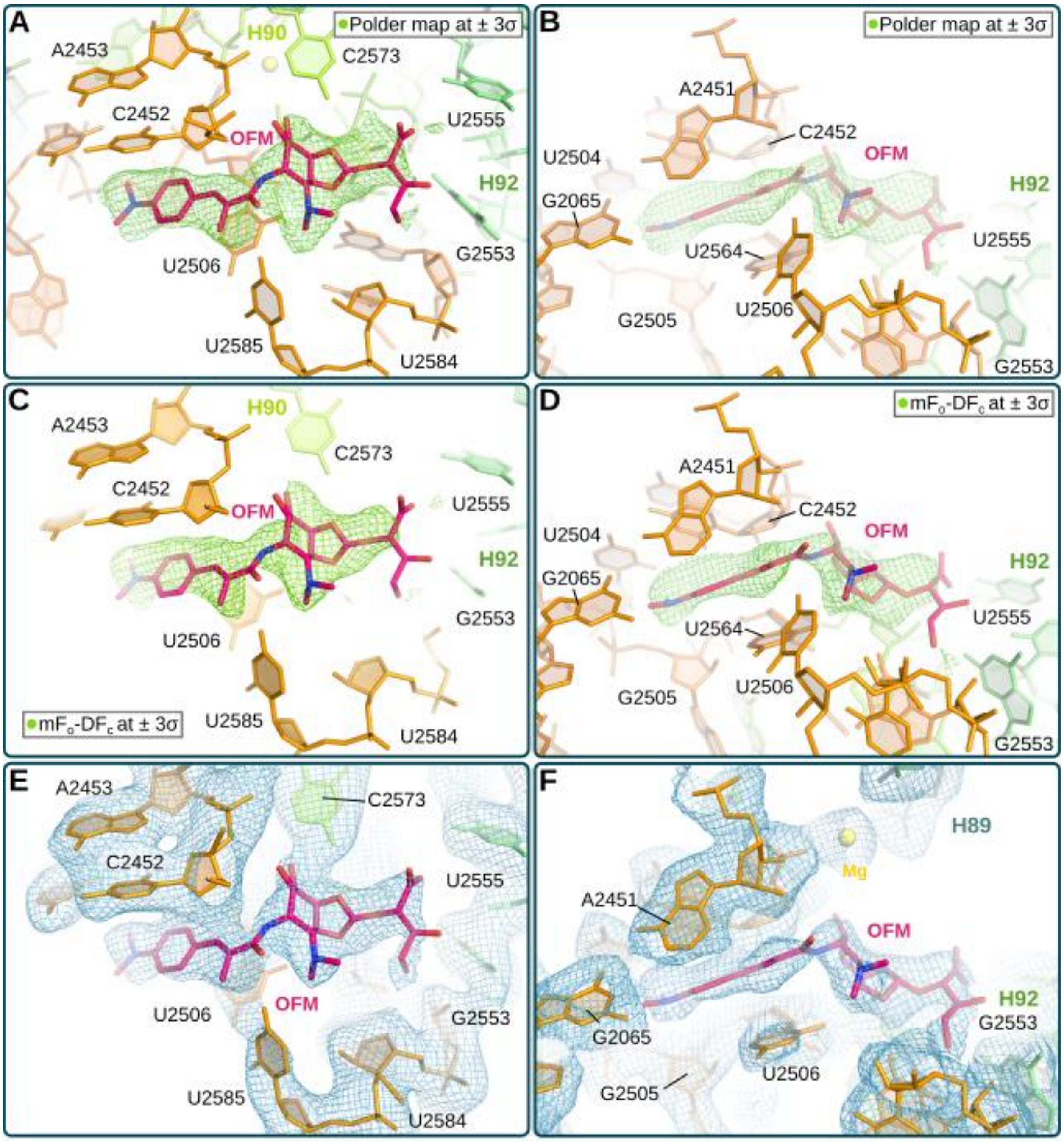


**Supplemental Figure 1: Electron density maps around the ligand binding site**. Top (**A**) and (**B**) side views of the initial (unbiased) polder map (*1*), which excludes bulk solvent around the area of interest, are shown at contour level of 3 σ. **(C-D)** The conventional mFo-DFc difference map is shown at a contour level of 3σ from the same views as the polder map. Although the terminal aromatic DMPE moiety [4-(dimethyl-amino)-phenyl]-2-methylprop-2-enoic acid] and its planarity is clearly accounted by the conventional mFo-DFc map the conformation of the aminocyclitol part and the those of the terminal L-erythronic acid can be more readily identified in the polder map. **(E-F)** The final 2mFo-Dfc map sharpened and contoured at 1 σ shows the ligand, highlighting in (**F**) the quasi planarity of the DMPE moiety with the conjugated peptide bond-like part.

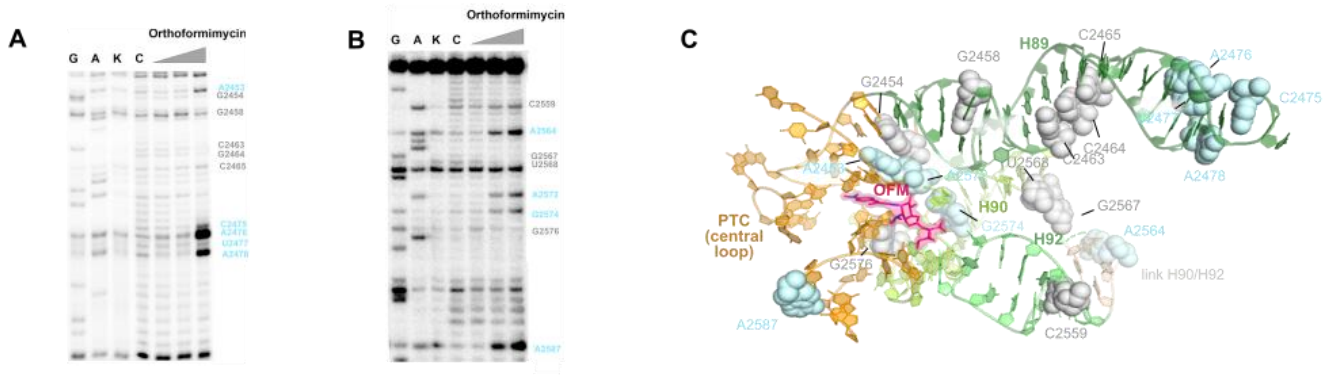


Supplemental Figure 2: The effect ofm on 23S rRNA accessibility to hydroxyl radical cleavage. Primer extension analysis was used to define the hydroxyl radical accessibility pattern of the (A) H89and (B) H92 in the absence (lane C) or presence of increasing amounts of ofm (1μM, 10 μM, and 100 μM; grey triangle). G and A are sequencing lanes and lane K contains the control 23S rRNA sample not subjected to cleavage. (C) Nucleotides that display decreased (grey) or increased (blue) accessibility are mapped onto the 50S-ofm structure.

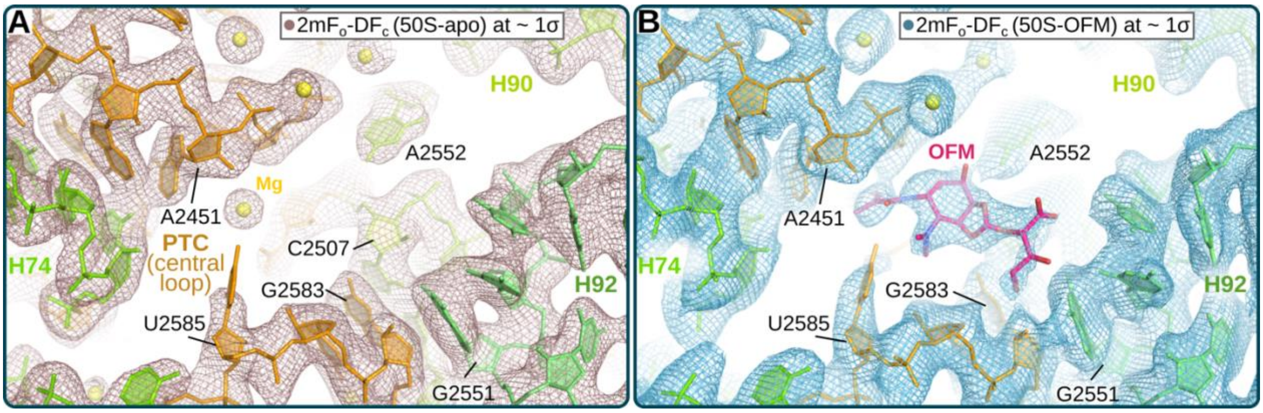


**Supplemental Figure 3: Final electron density maps for the 50S-apo and 50S-OFM structures**. Comparing the 2Fo-Fc maps from the (**A**) 50S-apo and (**B**) 50S-OFM structures highlights the displacement of a Mg ion in the ofm binding pocket and the significantly stronger density for the base of residue U2585 (due to ligand induced stabilization of the trans U2506-U2585 base interaction).
